# Impact of anatomic variability and other vascular factors on lamina cribrosa hypoxia

**DOI:** 10.1101/2024.09.12.610282

**Authors:** Yuankai Lu, Yi Hua, Po-Yi Lee, Andrew Theophanous, Shaharoz Tahir, Qi Tian, Ian A. Sigal

## Abstract

Insufficient oxygenation in the lamina cribrosa (LC) may contribute to axonal damage and glaucomatous vision loss. To understand the range of susceptibilities to glaucoma, we aimed to identify key factors influencing LC oxygenation and examine if these factors vary with anatomical differences between eyes. We reconstructed 3D, eye-specific LC vessel networks from histological sections of four healthy monkey eyes. For each network, we generated 125 models varying vessel radius, oxygen consumption rate, and arteriole perfusion pressure. Using hemodynamic and oxygen supply modeling, we predicted blood flow distribution and tissue oxygenation in the LC. ANOVA assessed the significance of each parameter. Our results showed that vessel radius had the greatest influence on LC oxygenation, followed by anatomical variations. Arteriole perfusion pressure and oxygen consumption rate were the third and fourth most influential factors, respectively. The LC regions are well perfused under baseline conditions. These findings highlight the importance of vessel radius and anatomical variation in LC oxygenation, providing insights into LC physiology and pathology. Pathologies affecting vessel radius may increase the risk of LC hypoxia, and anatomical variations could influence susceptibility. Conversely, increased oxygen consumption rates had minimal effects, suggesting that higher metabolic demands, such as those needed to maintain intracellular transport despite elevated intraocular pressure, have limited impact on LC oxygenation.

## 1. Introduction

Millions of people worldwide suffer from blindness or reduced vision due to glaucoma, a disease characterized by the degeneration of retinal ganglion cells and their axons.[1, 2] In glaucoma, axonal damage initiates at the lamina cribrosa (LC) region within the optic nerve head (ONH), compromising the transport of visual information from retina to the brain, and ultimately leading to vision loss.[3, 4] The LC receives blood with nutrients and oxygen through a complex and dense vascular network that is intertwined with the collagen beams and the retinal ganglion cell axons [5–9] Although the precise mechanisms of axon damage in LC remain unclear, one of the leading hypotheses posits that insufficient perfusion and oxygenation within the LC may contribute to cause the axonal damage.[10–15]

The LC microcirculation could be influenced by various factors, including vascular network geometry, perfusion blood pressure, tissue metabolic demands, tissue deformations, autoregulation responses, and tissue remodeling mechanisms.[6, 16–18] These factors could act independently or interact with each other, impacting both tissue perfusion and oxygenation, and thereby influencing physiological and pathological scenarios. Moreover, elevated intraocular pressure (IOP), one of the primary risk factors for glaucoma, could also contribute to this process. Elevated IOP can lead to deformation, compression, and distortion of the LC vasculature, compromising blood and oxygen supply to the LC region. [12, 17, 19–21] Our previous work also showed that LC oxygenation is more susceptible to systematic IOP-induced deformation than stochastic vasculature damage. [22] However, the critical threshold of IOP for glaucoma varies among patients and many individuals with elevated IOP never suffer full vision loss due to glaucoma.[23] To address the variation among different eyes and identify the most influential factors, a systematic analysis involving multiple eyes and various potential risk factors affecting LC hemodynamics and oxygenation is necessary. Such systematic investigations are crucial for advancing our understanding of glaucoma pathophysiology and developing more effective diagnostic and therapeutic strategies.

Despite major advances of imaging techniques over the last several years, such as optical coherence tomography angiography (OCT-A)[24–27], ultrasound techniques[28], MRI-based techniques[29], and laser speckle flowgraphy (LSFG)[30], obtaining direct measurements of LC hemodynamics and oxygenation with adequate resolutions and depth penetration remains elusive. Therefore, alternative approaches, such as theoretical modeling and numerical simulations, have been employed to assess blood flow and oxygenation within the LC region.[12, 13, 19, 22, 31, 32] Recently, we utilized experimentally-derived reconstructed 3D model of eye-specific LC vasculature and computational techniques for analyzing hemodynamics and oxygenation, and their influential factors. [32] We found that vessel radius, oxygen consumption rate, and arteriole perfusion pressure were the three most significant factors influencing LC oxygenation. However, the study was preliminary and based on a single eye anatomy. Eyes, however, vary in anatomy and a result applicable to one eye is not necessarily exactly the same for another. Hence, it remains unclear whether the factor influences identified on our previous study generalize to other eyes.

Our goal in this study was to identify the most influential factors and address the impact of eye anatomy differences on LC oxygenation. To achieve our objective, four eye-specific 3D models of the LC vasculature were reconstructed based on histological sections. From this, numerical simulations were performed to evaluate LC hemodynamics and oxygenation. Specifically, we parameterized the vessel radius, oxygen consumption rate, and arteriole perfusion pressure to evaluate their impact on LC oxygen supply. We used regularized grid to generate parameter combinations, allowing for the analysis of the independent and correlated effects of each parameter.

## 2. Methods

*General procedure*. First, we labeled, imaged, and reconstructed the eye-specific ONH vasculatures of four healthy monkeys as our baseline models, following the technique described elsewhere.[33, 34] Based on the four baseline vascular network models we then created 500 models by varying vessel radius, neural tissue oxygen consumption rate and arteriole blood pressure (four eyes, five levels per parameter, three parameters: 4 x 5^3^ = 500). Second, we performed hemodynamics and oxygenation simulations to evaluate the blood supply and from this the oxygen field within the ONHs. Third, we compared the fraction of hypoxic regions and the minimum oxygen tension within the LC across all models to determine the factor influences on LC hemodynamics and oxygenation. The steps are described in detail below.

### 2.1 Reconstruction of 3D eye-specific LC vascular network

All procedures were approved by the University of Pittsburgh’s Institutional Animal Care and Use Committee (IACUC), and followed both the guidelines set forth in the National Institute of Health’s Guide for the Care and Use of Laboratory Animals and the Association of Research in Vision and Ophthalmology (ARVO) statement for the use of animals in ophthalmic and vision research.

#### Vessel labeling

We processed four healthy female rhesus macaque monkeys’ heads for vessel labeling. Within two hours after sacrifice, we cannulated the anterior chamber of each eye to control IOP throughout the experiment. IOP was set to 5 mmHg to prevent hypotony or hypertension. Two polyimide microcatheters were inserted into the carotid arteries for labeling. Warm phosphate-buffered saline (PBS) was perfused to wash out intravascular blood first. A lipophilic carbocyanine dye, Dil, was then used to label the vessels in the eye. We perfused 100 ml Dil solution into each carotid artery, followed by a 100ml 10% formalin perfusion to fixate the eye.

#### Histology and 3D vasculature reconstruction

The ONH and surrounding sclera were then isolated, cryoprotected, and sectioned[35]. Each section underwent imaging through Fluorescence Microscopy for vessel visualization and Polarized Light Microscopy for collagen visualization.[33, 36] 3D vasculature reconstruction involved importing and registering sequential these images in Avizo software (version 9.1). Vessel segmentation aided by a Hessian filter, and vessels in the LC region were identified by the presence of collagen beams.[33] Our reconstructed vascular networks included the whole LC region, and some of the pre-laminar and retro-laminar regions (Figure 1). This ensured that the 3D LC network was fully enclosed within the region reconstructed.

**Figure 1.**
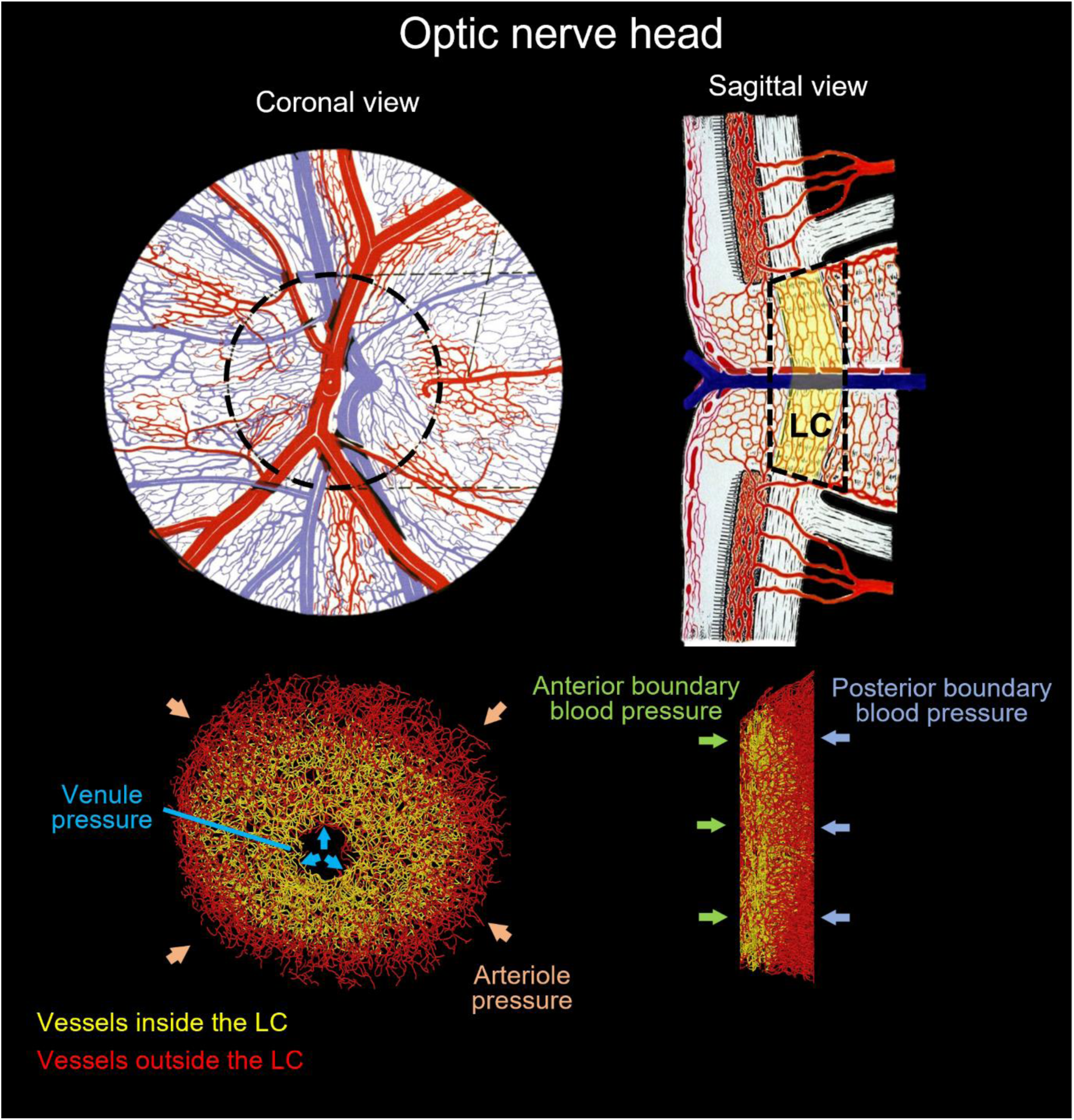
(Top) Diagram of the ONH adapted from [64]. Our model represents the vessels within the scleral canal, included the whole LC region, and some of the pre-laminar and retro-laminar regions. Black dashed lines represent the model boundaries, yellow area represents the LC region. (Bottom) An example eye-specific vessel network. To improve flow boundary conditions the region reconstructed extended beyond the LC. Vessels within the LC are shown colored in yellow. Vessels reconstructed but outside the LC are shown in red. The network is labeled to illustrate the blood pressure boundary condition settings. Four blood pressure conditions were assigned at the peripheral, central, anterior, and posterior boundaries of the model. See the main text for the rationale and details on how these pressures were assigned.

#### Vessel diameter setting

Although our reconstruction technique could be leveraged to obtain the vessel diameter information, we acknowledge that post-mortem changes, tissue swelling, and pressure variations can alter vessel diameters from in vivo conditions. Autoregulation, particularly in the deep optic nerve head (ONH) and lamina cribrosa (LC), remains uncharacterized in vivo. Following previous studies of LC hemodynamics [12, 13, 22, 31, 32], we assumed that all vessels in the LC had the same diameter. A uniform capillary radius of 4 µm was selected based on prior measurements[37], as discussed in [32]. Further research, combining our technique with in vivo imaging, could improve our understanding of vessel diameters and autoregulation in the LC.

#### Vascular reconstruction validation

The post-mortem dye perfusion process may encounter challenges, such as incomplete penetration into all vessel segments due to intravascular clotting or insufficient perfusate volume. To reduce clotting, we minimized the interval between animal death and perfusion and thoroughly washed the ONH with PBS. Our imaging revealed strong signals in retinal and choroidal vessels, indicating successful perfusion and sufficient labeling. Nevertheless, we acknowledge that vessel visualization and reconstruction could be impacted by uneven labeling, discontinuities, or leaks. Manual corrections, including ‘cleaning’ and ‘bridging’ segments, were performed to address these issues, though they may introduce artifacts and randomness.[32, 33] To evaluate the reconstruction technique, two students independently reconstructed the same labeled eye, and their results were compared to assess the method’s validity. The similarity in geometry and oxygenation observed in the two reconstructed networks demonstrated robustness in our approach, with minimal impact from artifacts introduced through manual correction.

### 2.2 Hemodynamics and Oxygenations

#### Oxygen diffusion in tissue

We followed the approach described in [38] to perform the LC hemodynamics and oxygenation simulations. Oxygen transport within the oxygen-consuming neural tissues can be described by the reaction-diffusion equation [39]:

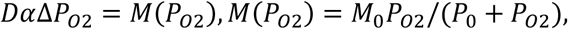

where D is the oxygen diffusion coefficient, α is the oxygen solubility coefficient, and *P_02_* is the tissue oxygen partial pressure. The oxygen consumption term *M*(*P_02_*) can be estimated by Michaelis-Menten enzyme kinetics [39], where *M*_0_ represents oxygen demand and *P*_0_ is the oxygen partial pressure at half-maximal consumption. In this study, *M*_0_ was assumed to be uniform throughout the LC.

#### Oxygen flux in blood

ONH vascular network system was considered as a set of interconnected capillary elements. The capillary elements connect with each other at the bifurcation nodes. The blood flow within a capillary can be approximately by the Poiseuille flow due to the low Reynolds number (< 0.05) [40],

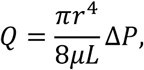

where Q is the flow rate, r is the vessel radius, L is the vessel length, *μ* is the blood viscosity, and Δ*P* is the pressure drop along the vessel.

Pressure boundary conditions were applied to the vascular network based on the physiological blood supply to the ONH. Specifically, we divided the model boundaries into four surfaces to specify the blood pressure conditions that supply the ONH region (see Figure 1). The inlet pressure for the arteriole, representing blood inflow from the circle of Zinn-Haller, was set to 50 mmHg at the periphery. The outlet pressure for the central retinal vein, responsible for drainage, was set to 15 mmHg. The anterior ONH boundary and the posterior ONH boundary were assigned pressures of 20 mmHg and 16 mmHg, respectively.

The oxygen flux in blood vessels satisfies:

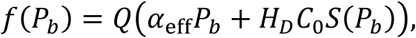

Where Q is the flow rate, *α*_eff_ is the effective solubility of oxygen in plasma, *H_D_* is the hematocrit, *C*_0_ is the concentration of hemoglobin-bound oxygen in a fully saturated red blood cell, *P_b_* is the blood oxygen partial pressure (mmHg), *S*(*P_b_*) is the oxygen-hemoglobin saturation, determined by the empirical oxygen–hemoglobin dissociation curve.

#### Oxygen exchange on vessel walls

Conservation of oxygen along each vessel segment implied that,

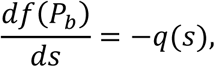

where *s* is arc-length parameter along the vessel, and *q*(*s*) is the total oxygen flux through the blood vessel wall per unit vessel length.

At the interface between blood vessel and tissue, the diffusive oxygen flux across the vessel wall must be consistent with the surrounding tissue oxygenation, implying that,

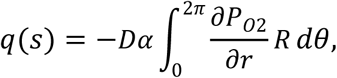

where *r* is the radial distance from the vessel centerline, R is the vessel radius, and the integral is around the circumference, denoted by angle *θ*. Therefore, the oxygen flux can be evaluated from the average gradient of *P_02_* on the vessel wall.

As reported before[38], we used a fast and efficient method to simulate the convective and diffusive oxygen transport in the complex ONH vascular networks. All parameters are listed in Table 1.

**Table 1.** Parameters used in hemodynamic and oxygenation simulations.

| **Constant parameters** | **Value** | **Reference** |
| --- | --- | --- |
| Oxygen diffusion coefficient, $D\alpha$ | $6 \times 10^{-10}$ mlO<sub>2</sub>/cm/s/mmHg | [39] |
| Effective oxygen solubility, $\alpha_{eff}$ | $3.1 \times 10^{-5}$ mlO<sub>2</sub>/ml/mmHg | [13, 39] |
| Oxygenation at half-maximal consumption, $P_0$ | 10.5 mmHg | [39] |
| Maximal RBC oxygen concentration $C_0$ | 0.5 mlO<sub>2</sub>/ml | [39] |
| Venule pressure | 15 mmHg | [32] |
| Anterior blood pressure | 20 mmHg | [32] |
| Posterior blood pressure | 16 mmHg | [32] |
| **Parametric factors** |  |  |
| Vessel radius | 4 μm | [32] |
| Arteriole pressure | 50 mmHg | [32] |
| Consumption rate, $M_0$ | $5 \times 10^{-4}$ mlO<sub>2</sub>/ml/s | [32] |

### 2.3 Parametric analysis

We performed a parametric analysis based on the variation of three parameters: vessel radius, neural tissue oxygen consumption rate, and arteriole pressures. These parameters were selected because they had the strongest influences on LC hemodynamics and oxygenation in our previous study.[32] Baseline values of the parameters were established from the literature, as shown in Table 1 and discussed in detail [32]. To provide a systematic and unbiased parametric analysis these were all varied ±20% of the baseline (80% to 120%). Five parameter levels were selected: 80% (low), 90%, 100% (Baseline), 110% and 120% (high), resulting in a total of 5*5*5=125 models for each eye. We repeated the same parametric analysis for all four eyes, resulting in a total of 500 models.

As outcome measures, we used the minimal oxygen tension in the LC and the tissue volume fraction of the hypoxia region [41–43]. For minimal oxygen tension, the 10^th^ percentile was used as the definition of the minimal value to reduce the influence of very small regions that could be artifactual [32]. For the tissue volume fraction of hypoxia, we defined hypoxia region as the region where oxygenation is lower than 38 mmHg (5% saturation), as in our previous work [22].

### 2.4 Statistical analysis

We utilized ANOVA to assess the rank and statistical significance of all parameters. Specifically, we used the percentage of the total sum of squares corrected by the mean as a metric to represent the contribution of each parameter and interaction.[15] The results among different eyes were analysis collectively, where the eye itself was considered as a categorial factor in ANOVA to account for individual variations in our models. Further details regarding the statistical analysis can be found in our previous parametric work.[32]

## 3. Results

The baseline hemodynamics and oxygenation of the four networks are shown in Figure 2. Although the eyes differed in anatomy, their hemodynamic and oxygenation exhibited similar characteristics. The flow rate and oxygenation were high at the periphery and low at center. This pattern is consistent with the blood supply for LC, where blood perfusion is from the periphery, draining through the central retinal vein. Detailed illustrations of flow and oxygen pattern are shown in Figure 3 and Video 1 (See Supplemental Material).

**Figure 2.**
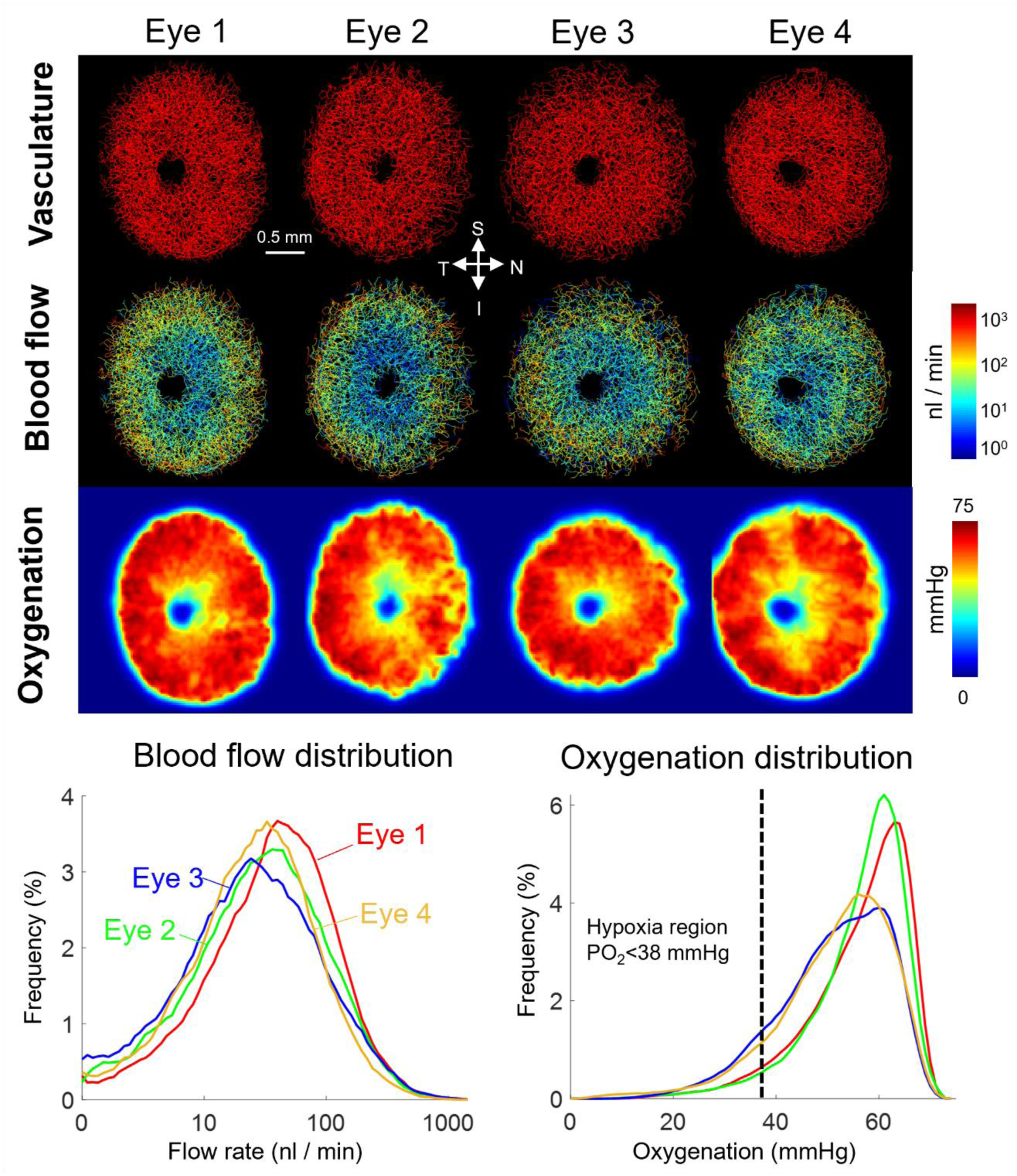
Vascular geometry, baseline hemodynamics, and baseline oxygenation of four eyes. Top: Vascular geometry and maps of blood flow and oxygenation. Bottom: Distributions of blood flow and oxygenation in the LC region. Blood flow and oxygenation exhibited similar features and tendencies across all eyes. Higher flow rates and oxygenation occurred at the periphery of the LC region, and lower flow rates and oxygenation were observed at the center. Interestingly, the distribution curves of blood flow and oxygenation showed different patterns. Blood flow exhibited significant variation across the entire LC, ranging from 1 to 1000 nl/min, while the LC was consistently well-provided with oxygen. Oxygen tensions remain consistently high throughout most of the LC region, with minimal regions experiencing hypoxic conditions.

**Figure 3.**
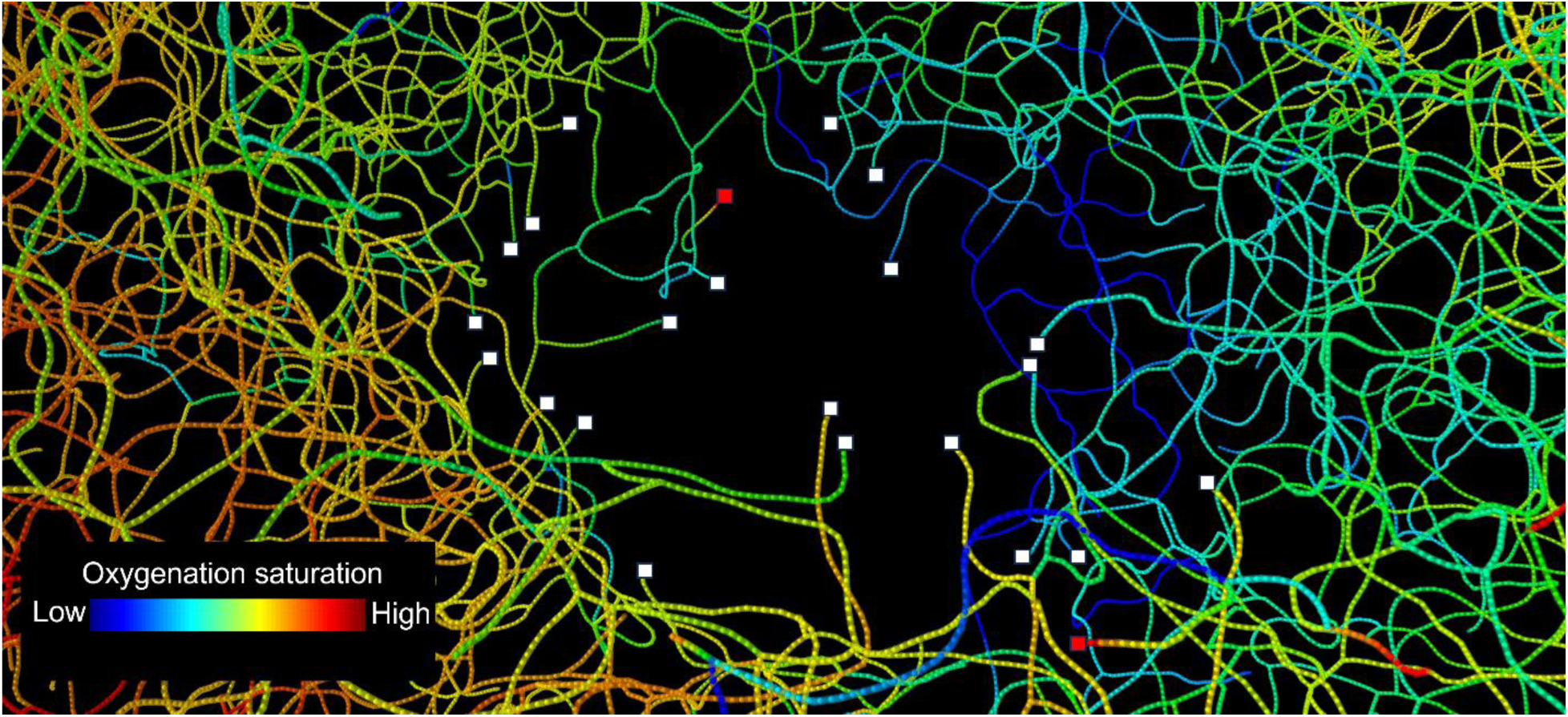
A screenshot of the video illustrating red blood cell transport in the blood vessels. The region shown is a close-up of the drainage at the central retinal vessels. Colors represent the blood oxygen saturation. The dots along each vessel represent the red blood cells. White and red squares represent blood flow outlets and inlets, respectively. The vessel ends in the center of the LC serve as flow outlets, according to the boundary conditions of flow drainage through the center retinal vein. Vessel ends that are not draining are because they end at the model anterior/posterior boundary (see Figure 1) and thus flow can be to/from the LC. Notably, the blood oxygen saturation exhibits an asymmetric pattern at the center, with lower oxygenation observed on the right side (Nasal side).

Figure 4 illustrates oxygenation distribution under various radii, consumption rates, and arteriole pressures for Eye 1. We selected three levels—low (80%), baseline (100%), and high (120%)— to show the impact of each parameter on LC oxygenation. Notably, changes in vessel radius caused the largest changes in LC oxygenation distribution, while variations in other parameters had comparatively smaller effects.

**Figure 4.**
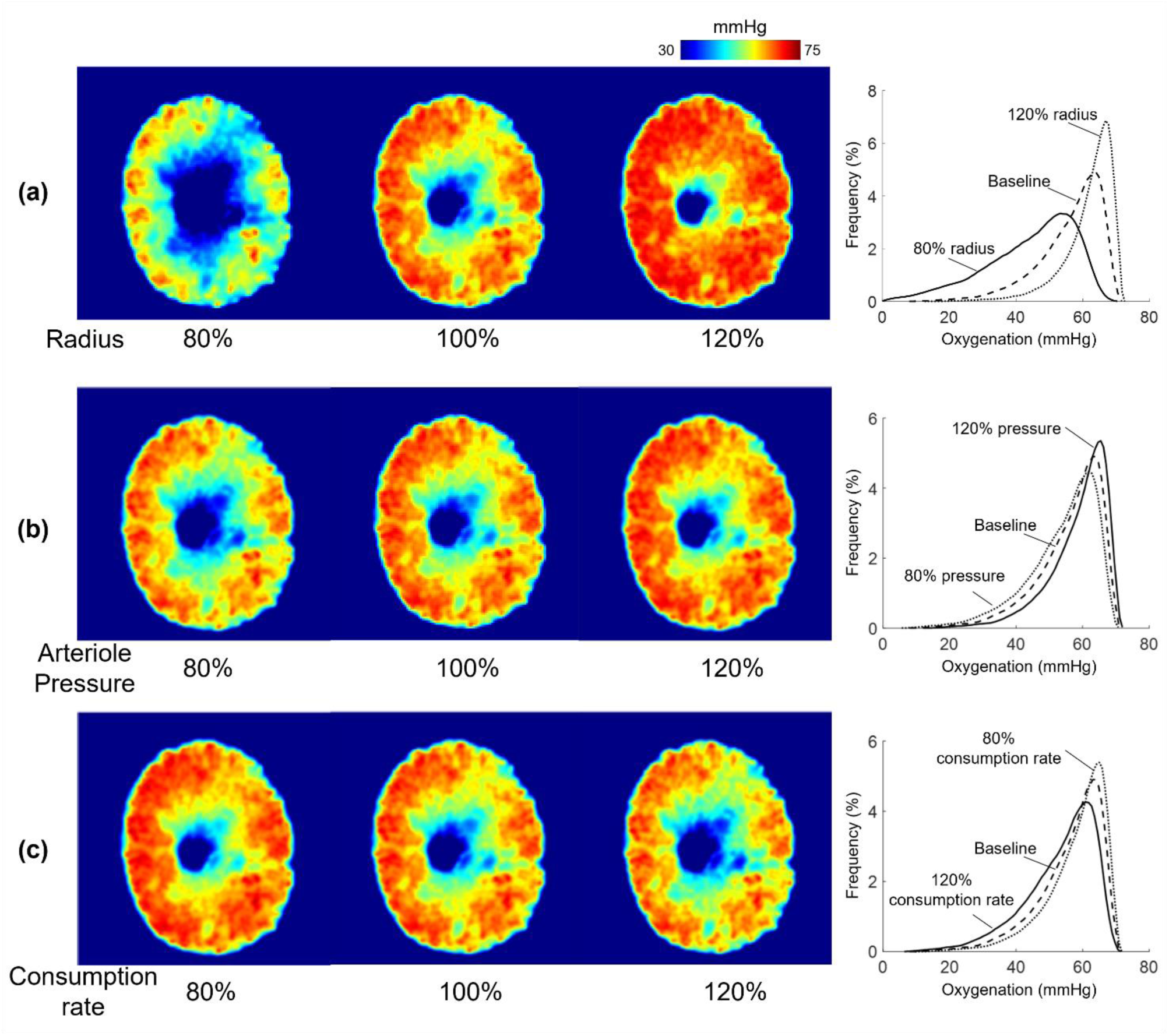
Oxygenation distribution across various radii, consumption rates, and arteriole pressures for eye 1. (a) Oxygenation at different radii. (b) Oxygenation at different Arteriole pressures. (c) Oxygenation at different consumption rates. Three levels—low (80%), baseline (100%), and high (120%)—were selected to illustrate the impact of each parameter. Radius demonstrated the strongest positive influence on oxygenation, while consumption rate and arteriole pressure had minor effects. Most hypoxia regions were observed near the center of the LC across all parametric scenarios.

Boxplots of the impact of each factor on the 10th percentile oxygenation and the fraction of hypoxic regions across all eyes are presented in Figure 5. Vessel radius exhibited the strongest positive relationship with the 10th percentile oxygenation and the strongest negative relationship with the fraction of hypoxic regions, consistent with the results shown in Figure 4. The statistical significance of each parameter was evaluated via ANOVA (see Figure 6). The most influential factors were vessel radius, arteriole pressure, and oxygen consumption rate. There were week interactions between these influential factors. Specifically, the interactions between radius and pressure are illustrated in Figure 7.

**Figure 5.**
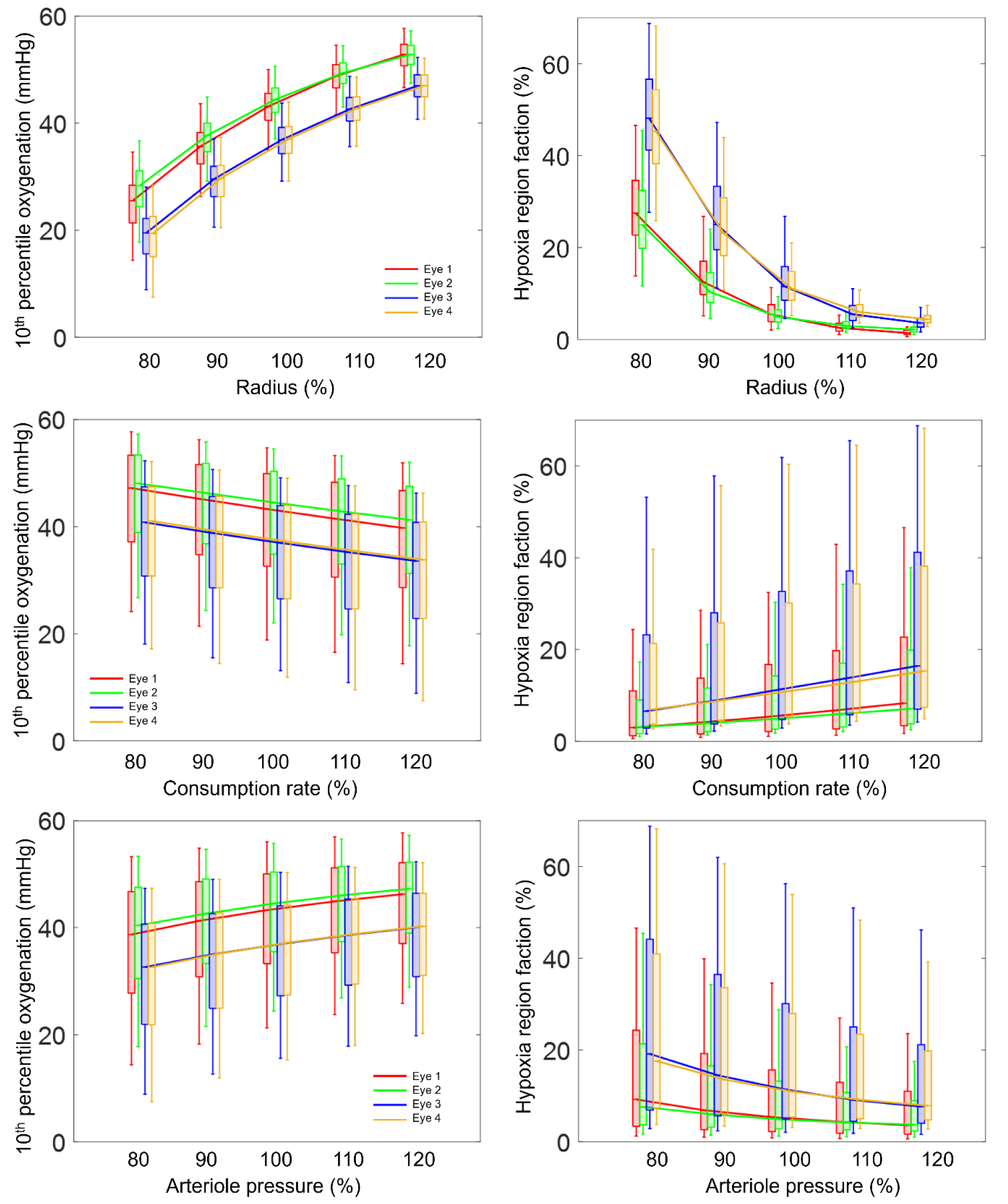
Boxplots showing the factor influences on the 10^th^ percentile oxygenation and hypoxia region fraction in the lamina cribrosa across all eyes. The top and bottom edges of each box are the upper and lower quartiles (25^th^ and 75^th^ percentiles), while the line inside of each box is the sample median (50^th^ percentiles), respectively. The end of the whiskers shows the minimum and maximum. To aid visualization we added lines connecting the median values. Vessel radius shows the strongest positive relationship with the 10th percentile oxygenation and the strongest negative relationship with the hypoxia region fraction. The variation observed among different eyes exceeds the oxygenation difference between scenarios with 80% consumption rate/arteriole pressure and those with 120% consumption rate/arteriole pressure. Interestingly, Eyes 1 and 2 exhibited similar LC oxygenation across all parametric cases, while Eyes 3 and 4 also showed similar patterns to each other but differed from Eyes 1 and 2. The changes in LC oxygenation in response to the parameters, or in other words, the sensitivity to parameters, were consistent across all eyes.

**Figure 6.**
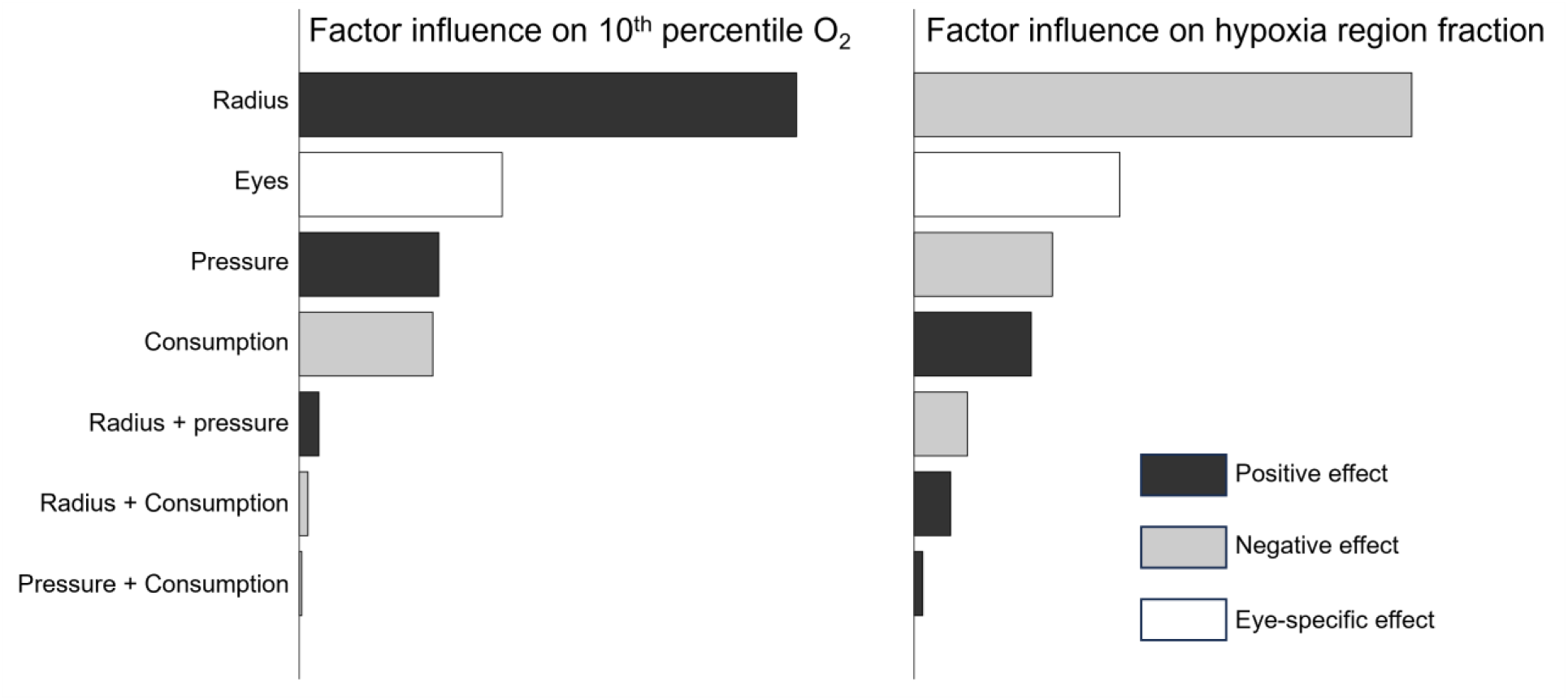
Bar chart showing the ranking of factors and interactions according to their influence on the 10^th^ percentile oxygenation and hypoxia region fraction in the LC, as determined by ANOVA. The vessel radius was strongest influence factor, followed by the eye, arteriole pressure, and consumption rate. Interactions between the parameters show minor contributions to the LC oxygenation.

**Figure 7.**
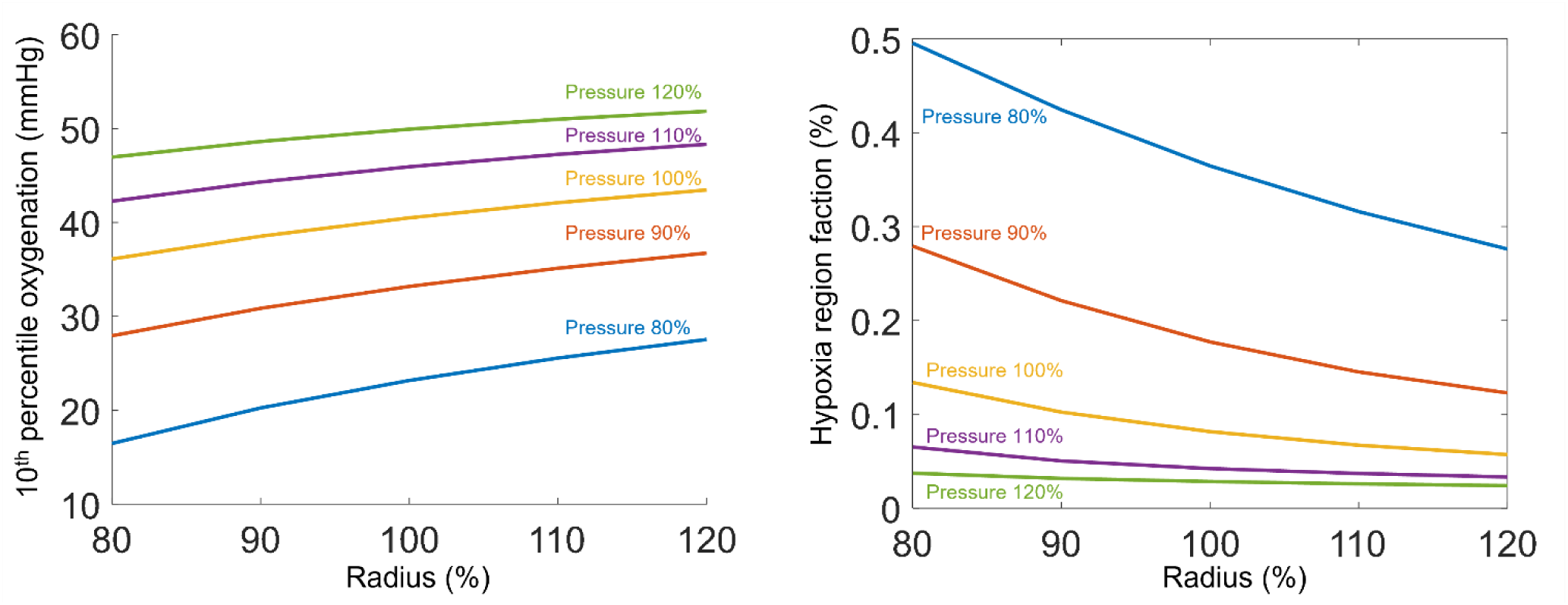
Interactions between vessel radius and arteriole pressure on LC oxygenation. Regarding the 10^th^ percentile oxygenation, the curves for different pressures were nearly parallel, suggesting a weak interaction between the radius and pressure parameters. For hypoxia region fraction, the influence of the pressure was stronger when the radius was low.

## 4. Discussion

Our goal was to identify the most influential factors on LC oxygenation, including the impact of eye to eye anatomical variations. Specifically, we used four eye-specific 3D LC vasculature models and parameterized the vessel radius, oxygen consumption rate, and arteriole perfusion pressure to evaluate their impact on LC oxygen levels. The four most influential factors on LC oxygenation were, from most to least influential: vessel radius, eye anatomy, arteriole perfusion pressure, and oxygen consumption rate. Our models also showed that the LC was well irrigated at baseline IOP. Below we discuss the findings in more detail as well as the limitations and other considerations to keep in mind when interpreting them.

### Vessel radius was the most influential factor on the LC oxygenation

Our models predicted that the vessel radius has the strongest contribution to LC oxygenation, both for the 10^th^ percentile oxygenation and hypoxia region fraction. This can be understood as follows: Vessel radius plays a crucial role in determining the flow resistance of each vessel segment, as evidence by the well-known quartic power in Poiseuille’s flow formula. By precisely controlling luminal radius through vessel wall contraction and dilation, this sensitivity can be leveraged by vascular systems to regulate blood and oxygen supply efficiently. There are, however, some scenarios in which the high sensitivity to vessel radius could prove problematic to LC perfusion. For example, IOP-induced deformations can distort the LC tissues, altering the geometry of the vessels within, altering radius. Substantial experimental evidence supports the idea that blood perfusion in the ONH decreases and vessel radius reduces with elevated IOP.[13, 31, 44–46] Thus, our model predictions are consistent with the literature. Given the complexity of the LC mechanics and vascular network, predicting the effects of IOP-related deformations on LC oxygenation is not trivial. Elsewhere we have presented a combined experimental-computational analysis using models similar to those in this work.[47] We found that moderately elevated IOP can cause sufficient distortions to the LC vasculature to alter LC hemodynamics and lead to mild hypoxia in a substantial part of the LC. More extremely IOP elevations can lead to severe hypoxia that is likely to cause more immediate damage to the LC neural tissues.

### Eye anatomy variations had the second strongest influence on the LC oxygenation

Eye anatomy variations refer to the differences in oxygenation between eyes. This means that for vascular networks modeled with exactly the same parameters and differing only on the vascular network, the hemodynamics and oxygenation were substantially and significantly different. As shown in the Results section, the four models formed two groups. See, for example, the plot of LC oxygenation in Figure 2. The models of Eyes 1 and 2 show similar curves, distinct from the curves of eyes 2 and 4. It is important for readers to recall that the response curves will change as other parameters vary, however, the ANOVA results indicate that the changes in the curves will be similar for all the eyes since there were no significant interactions between eye anatomy and other parameters. This means that eyes differ in hemodynamics and oxygenation, but that their sensitivity to changes in other parameters is the same. Thus, while it is still not clear why some eyes, or their vascular networks, behave differently, there is a common similar sensitivity. Some eyes may be more susceptible to low oxygenation than others, but it seems to be that this is not because of a higher sensitivity to the parameters. Our recent work investigated the IOP effect on LC oxygenation across different eyes. [47] Different eyes anatomy led to different LC oxygenation. However, the changes of LC oxygenation due to IOP-induced deformation were similar for all eyes, which also suggest the common sensitivity.

Previously we conducted a study similar to this one, using a 3D eye-specific model of the LC vasculature to evaluate how hemodynamics and oxygenation were affected by varying several parameters. [32] We found that vessel radius, oxygen consumption rate, and arteriole perfusion pressure were the three most significant factors influencing LC oxygenation. In that study, however, we used only one vascular network and therefore were unable to evaluate the effects of eye anatomy. This is why it was crucial to conduct the study described in this paper.

Other studies have looked at variations in LC hemodynamics resulting from differences or changes in anatomy. Most of these studies have considered the LC as a generic model, varying parameters such as the LC size, depth and curvature.[12, 13, 31, 48] These studies found that several structural parameters of the LC, such as LC diameter and cup depth, had a more stronger influence on LC oxygenation than perfusion pressure. Other structural parameters, like anisotropy and pole size, had a weaker impact on LC oxygenation. Overall, LC structural parameters, or eye anatomy, play a crucial role in influencing LC oxygenation, which is consistent with our findings in this work.

The simplified generic models have several important strengths, as we have discussed elsewhere [49], but there are also disadvantages. Generic models can potentially miss core details of the architecture of a specimen. Also, it is possible that the parameter space considered by the generic models does not represent the actual variability of eyes, not just in terms of the parameters and their ranges, but their distributions. If this is the case, the sensitivities may be inaccurate. Further, it is possible that generic models fail to account properly for model complexity and variability and thus miss important conditions. This is where the use of eye-specific models reveals an important strength that we leveraged in this work. Eye anatomy, however, remains a discrete parameter. Elsewhere we have shown techniques that potentially could be adapted to parameterize eye-specific models. [32]

### Arteriole perfusion pressure and oxygen consumption rate ranked third and fourth most influential factors on LC oxygenation

Arteriole perfusion pressure and oxygen consumption rate ranked as the third and fourth influential factors on LC oxygenation. The LC oxygenation was positively associated with the arteriole perfusion pressure, and negatively associated with the oxygen consumption rate. LC oxygenation reflects the balance between supply and consumption. Increased perfusion pressure would drive more blood flow across the LC and bring more oxygen to neural tissues. Conversely, a higher oxygen consumption rate would consume more oxygen in LC, potentially leading to hypoxia under insufficient supply. Perfusion pressure came from the cardiovascular circulation, and was affected by the upstream vascular systems and some blood flow regulation mechanism. Cardiovascular dysfunctions, such as hypo/hypertension, were found to be linked to glaucoma development.[50–53] It is worth noting that the hypo/hypertension will also induce the remodeling of vascular structures, such as systemic vasoconstriction (vessel radius reduction), capillary rarefaction, which may also alter the LC hemodynamics and oxygenations.[54–56] The oxygen consumption rate in LC varied based on the amount, type, and activity of various cells in neural tissues, including axons, astrocytes, and other cells, and could even change during the glaucoma development. [43, 57]

From a perspective of physiological mechanisms and biological functions, the perfusion pressure and consumption rate should directly affect the LC oxygenation, which aligns with our expectations and findings. Interestingly, they contribute less than the eye anatomy variations.

### The LC was well irrigated at baseline

All the eyes were well irrigated under baseline conditions, with less than 15% of the LC volume suffering low enough oxygenation to be at risk of hypoxia. This is consistent with the idea that healthy individuals have low risk of LC hypoxia. The result also agrees with our previous work on the robustness of the LC oxygenation. [22]

To the best of our knowledge, this is the first study to conduct parametric analysis based on multiple 3D eye-specific LC models while accounting for eye anatomy. We combined the experimental-based reconstructed LC vasculature with highly accurate and efficient numerical simulation techniques to provide a high-resolution estimation of LC hemodynamics and oxygenation. Our reconstruction technique enables the creation of multiple LC vascular networks from histological sections of healthy monkeys, offering high-resolution and accurate structural detail. Utilizing a fast algorithm for hemodynamic and oxygenation simulations, we conducted a systematic parametric analysis across multiple eyes.

## Limitations

It is important to acknowledge the limitations of this study so that readers can consider them when interpreting our findings. The first limitation is that we used computational modeling. While our techniques have been tested and verified, much better experimental data is required to consider the predictions from our models confirmed and validated. The LC is extremely complex and there are still too many aspects that are not fully understood. Unfortunately, at this time, the experimental tools and techniques necessary to gather the data necessary do not exist. As discussed before, the challenges in spatial and temporal resolution and in signal penetration are not yet overcome. There are promising tools in the horizon, like visible light OCT that may enable measuring blood oxygenation [26] and super-resolution ultrasound that may allow measuring of blood flow with high resolution and deep into the ONH. [58] Also, while this study expanded on our previous parametric analysis of LC hemodynamics and oxygenation by including several eye-specific models, four models is not enough to capture the full variability in human or monkey eyes. Studies with more eyes, especially including human eyes, are crucial.

Although we reconstructed the 3D eye-specific LC vasculature model from experimental images, we recognize that there were gaps between our reconstructed vasculature model and the in-vivo LC vascular network. A salient one is that we set the vessel radius in LC as a uniform value. Currently, there are no studies offering detailed distributions of vessel radius in the LC, and the use of a uniform radius has been common in previous LC hemodynamic studies. Our reconstruction technique was based on ex-vivo histological sections. The difference in the internal environment (e.g., absence of blood pressure, interstitial fluid pressure, IOP) result in the different vessel radius from the in-vivo case. Since our analysis indicates that the vessel radius was the strongest influential factor for LC oxygenation, it is important to consider the limitation of our uniform radius assumption and the potential effect due to the non-uniform radius distribution. Further research, potentially with a more advanced reconstruction technique coupled with in vivo imaging, could help provide detailed information on in-vivo LC vessel radius and estimate the effect of the radius distribution.

Another limitation is that our work only considers the static status of LC hemodynamics and oxygenation. All the parameters in our model were assumed to be time-invariant and independent. However, in-vivo blood/oxygen supply involves several regulation mechanisms to meet the changing demand of organisms. For instance, short-term blood flow autoregulation has been identified as a significant factor in hemodynamics and oxygenation for eye.[16, 59] Long-term vessel remodeling in the LC has also been reported in the development of glaucoma.[2, 6] Although the precise regulation and remodeling in LC remain unknown, we acknowledge that they could alter the LC blood and oxygen supply.[8, 16, 18, 59–61] Future research should incorporate the dynamic aspects of the LC blood and oxygen supply, which might contribute to the development of pathology, such as glaucoma.

It is important to note that we did not consider IOP-induced vessel deformations in this study. Although IOP is a primary risk factors for glaucoma, the impact of IOP-induced deformation on LC oxygenation remains unknown. Elevated IOP may affect LC blood/oxygen supply directly by distorting LC capillaries, and/or indirectly by reducing perfusion pressure through the feeding vessels, e.g., posterior ciliary arteries (PCAs).[5, 62, 63] The effect of elevated IOP is complex and is therefore addressed in a dedicated study.[47]

In summary, we used four reconstructed eye-specific 3D LC vasculature, and parameterized the vessel radius, oxygen consumption rate, and arteriole perfusion pressure in our hemodynamic models to evaluate their impact on LC oxygen supply. Our model predicted that the vessel radius and eye variation had the most significant influence on the LC oxygen supply. Situations that alter the radius, such as IOP-induced deformation, may contribute to compromised LC oxygenation. But different individuals could also exhibit different susceptibility to those pathological scenarios due to their anatomy differences.

## Acknowledgements

National Institutes of Health R01-EY023966, R01-EY031708, R01-HD083383, P30-EY008098 and T32-EY017271 (Bethesda, MD), the Eye and Ear Foundation (Pittsburgh, PA), Research to Prevent Blindness (unrestricted grant to UPMC ophthalmology, and Stein innovation award to Sigal IA) and BrightFocus Foundation.

